# Structural clustering and functional profiling of NMAN-causing variants in HINT1 suggest personalized therapeutic strategies

**DOI:** 10.1101/2023.12.01.569336

**Authors:** Silvia Amor-Barris, Tamas Lazar, Ayse Candayan, Luiza L. P. Ramos, Kristien Peeters, Shoshana J. Wodak, Albena Jordanova

## Abstract

**Background:** Biallelic loss-of-function variants in *HINT1* cause neuromyotonia-associated axonal neuropathy (NMAN). Affected patients present from an early onset with a motor-greater-than-sensory polyneuropathy that is currently incurable. NMAN is a global cause of inherited peripheral neuropathy with higher prevalence in Europe and Asia. Nearly 30 distinct NMAN-associated variants have been reported to date, mostly in sporadic patients and small families. There is limited functional characterization of most of them, resulting in limited knowledge on their pathogenic mode of action and hindering the development of therapeutical strategies.

**Methods:** We systematically (re-)evaluated the pathogenicity of all reported *HINT1* variants associated with NMAN using several *in silico* pathogenicity predictors used in the standard of care genetic testing. Fifteen missense and one truncating variants were further mapped onto the HINT1 crystal structure and grouped according to their spatial distribution. We combined several structural modeling tools to assess the impact of the variants on protein stability in both monomeric and dimeric forms. These variants underwent further functional validation by immunoblotting in both HeLa *HINT1* knockout cell lines and in a *S. cerevisiae* strain deficient for the HINT1 orthologue. Finally, we tested the *in vivo* functionality of each variant by its ability to rescue yeast growth under stress conditions.

**Results:** Our combinatorial approach allowed the systematic characterization and (re-)evaluation of all known *HINT1* variants associated with NMAN, enabling improved pathogenicity classification. Additionally, this standardized functional assessment allowed their categorization into four structure-based groups: 1) truncating variants; 2) variants of the catalytic pocket; 3) variants at the dimer interface; 4) variants in the distal β-hairpin and nearby loop. Functional tests in yeast and mammalian disease models demonstrated their detrimental- yet differential -effect on HINT1 function, enabling structure-function correlations.

**Conclusions:** Our standardized experimental pipeline allows for the characterization of newly discovered HINT1 variants in the context of NMAN, facilitating their pathogenicity interpretation. This work highlights the value of combining structural and functional approaches to understand better the underlying disease mechanisms. Our findings provide the rational basis for patient stratification and the development of personalized treatment strategies.

## INTRODUCTION

The widespread adoption of massive parallel sequencing technologies dramatically increased the number of known disease-causing genes and variants [1–3]. Rare diseases have particularly benefited from this advancement and more than 180 novel disease-causing genes were identified within three years after the commercialization of whole-exome sequencing [3]. While these technological improvements have ended the diagnostic odyssey for many patients, they have yet to translate into widely available therapies. Over 6,000 unique rare diseases affecting an estimated 350 million people worldwide have been identified so far, with 80% having a genetic origin. Yet, there is an FDA-approved treatment available for only 5% of them[4]. In rare neuromuscular disorders, most available therapies focus on symptomatic relief or palliation, with no existing disease-modifying therapies or cures. Identifying novel treatment opportunities is challenging due to the poor understanding of underlying disease mechanisms, the small size of patient cohorts, and the limited commercial incentive for companies to invest in the costly development of such therapies [5, 6].

These challenges are exemplified in the case of neuromyotonia-associated axonal neuropathy (NMAN [MIM:137200], also known as HINT1 neuropathy). NMAN is an autosomal recessive disorder caused by the loss of functional histidine triad nucleotide-binding protein 1 (HINT1) [7]. Patients display a motor greater than sensory peripheral polyneuropathy with an age of onset mainly within the first decade of life. The disease is characterized by wasting and weakness of the distal limb muscles, sensory loss, and skeletal deformities. This slowly progressive chronic disease is currently incurable. A characteristic feature of HINT1 neuropathy is neuromyotonia, which is present in 70% of the patients, clinically manifests as delayed muscle relaxation after a voluntary contraction, and is caused by the hyperexcitability of the peripheral nerves [7, 8]. NMAN is reported in patients worldwide, yet it has a non-random geographic distribution. The neuropathy is most prevalent in Europe and Asia due to several founder variants (p.Arg37Pro, p.Cys84Arg, p.Gly89Val, p.Arg95Gln, p.Glu100Gly, p.His112Asn) [8]. Among those, p.Arg37Pro shows a high carrier frequency (1:67-1:250) in Central-Eastern Europe, contributing to the significant morbidity of NMAN in this part of the world [7, 9, 10].

HINT1 protein is part of the histidine triad (HIT) superfamily characterized by a highly conserved amino acid motif (His-X-Hix-X-His-X-X, where X= hydrophobic amino acid) in the catalytic core near the carboxyl terminus. It forms a homodimer that functions enzymatically as a purine phosphoramidase [11–13] and has also been described as a (de)SUMOylase [14]. *In vitro,* HINT1 hydrolyzes many different substrates, however, the endogenous substrate(s) and the biological relevance of these activities remain unclear. *In vivo*, HINT1 is a transcriptional regulator involved in apoptosis [15, 16], tumor suppression [17, 18], and cardiac hypertrophy [19]. In the central nervous system, it affects the endocannabinoid pathway by modulating the NMDAR activity [20–22]. In the peripheral nervous system, where its deficiency causes NMAN, its role remains utterly uncharacterized.

Nearly 30 biallelic variants have been causally associated with NMAN, most of which occur in sporadic patients and often in compound heterozygous states. Characterization of their impact on HINT1 activity was done in only a few cases. In this study, we performed a systematic structural and experimental (re-) evaluation of the impact of 16 selected NMAN-associated variants on HINT1 function, adding novel evidence for their pathogenicity. Importantly, we were able to group the variants based on their differential effect on the HINT1 protein structure and function. Our findings provide novel insights into the pathomechanisms behind NMAN, justifying the implementation of patient stratification strategies and the development of personalized treatment options.

## MATERIAL AND METHODS

### Selection of HINT1 variants

We conducted a comprehensive literature search using PubMed to identify HINT1 variants associated with NMAN that have been reported in peer-reviewed articles published between 2012 and 2024. For each reported variant, we determined allele frequency using data from the Genome Aggregation Database (gnomAD v4.1). Subsequently, we performed *in silico* analyses using pathogenicity predictions tools routinely employed in diagnostic settings, including CADD (v3) [23], REVEL (v4) [24], AlphaMissense [25], and SpliceAI [26].

### Solvent accessible surface area and in silico protein fold stability calculations

The effect of the missense variant on the protein stability was calculated using FoldX-4 [27], SDM [28], mCSM [29], DUET [30], ENCoM [31], and DynaMut [32]. These methods combine information from protein sequences, 3D structures, and crude physicochemical parameters to calculate the change in protein folding free energy (stability) (ΔΔG). The calculations were carried out on the isolated HINT wild-type monomer and on the homodimer to evaluate separately the effect of the variants on the stability of individual subunits and the dimer.

We calculated the solvent-accessible surface area of individual residues (ASA_R_) in the monomer and the dimer. Based on these, we calculated the buried surface area of a residue (BSA_R_) upon dimer formation representing the contribution of the residue to the dimer stability [33]; which fraction gets buried in the dimer (fBSA_R_= BSA_R_/ ASA_R_(monomer)), and the residue relative accessible surface area RSA_R_ = [ASA_R_ (monomer or dimer)/ ASA_R_(Gly-X-Gly)], where ‘x’ represents the same residue in the free dipeptide unit. The ASA_R_ of the residue in the ‘x’ position of this very flexible dipeptide is taken to represent its ASA in the unfolded state of the protein. The various solvent-accessible surface area calculations were performed with the software FreeSASA 2.0.3[34] using default parameters.

### Cloning and creation of HINT1 expression plasmids

Yeast and mammalian expression plasmids with human *HINT1* were generated in previous studies[7, 35]. *HINT1* variants were introduced with site-directed mutagenesis using the KAPA HiFi kit (Roche Diagnostics, Basel, CH). Verification of the variants was done using Sanger sequencing.

### Cell line establishment and culture

*HINT1* HeLa KO cell lines were created using CRISPR/Cas9, as described previously[35]. HeLa cells were maintained in high-glucose DMEM (Gibco, Waltham, MA, USA) supplemented with 10% heat-inactivated fetal bovine serum (FBS, Gibco, Waltham, MA, USA), 1% penicillin/streptomycin (Gibco, Waltham, MA, USA), and 1% glutamine (Gibco, Waltham, MA, USA). Cells were grown at 37°C and 5% CO_2_ in a humidified atmosphere.

### Cell transfection

HeLa cells were transiently transfected using polyethylenimine (PEI) MW25000 (Polysciences, Warrington, PA, USA) with a pCAGGAS vector carrying human WT or variant HINT1. A pCAGGS-MBP was used as an empty vector control. Cells were seeded in a 6-well plate the day before the transfection in a complete medium without antibiotics. Cells were transfected with PEI and 250ng plasmid when 70-80% confluency was reached.

### Immunoblotting

Cells were lysed in RIPA lysis buffer (20 mM Tris-HCl pH=7.4; 150 mM NaCl; 0.1% Nonidet P-40; 0.5% sodium deoxycholate; 0.1% sodium dodecyl sulfate) supplemented with Halt™ Protease Inhibitor Cocktail (ThermoFisher Scientific, Waltham, MA, USA) for 30 min on ice and cleared by centrifugation for 10 min at 14000 RPM. Protein concentration was measured with the Pierce BCA protein assay kit (ThermoFisher Scientific, Waltham, MA, USA) and adjusted to 20µg per sample. Lysates were boiled for five min at 95°C in reducing Laemmli sample buffer (Bio-Rad, Hercules, CA, USA) supplemented with 100mM 1.4-Dithiothreitol (DTT).

Yeast proteins were extracted following a previously published protocol[35]. Briefly, yeast cells were collected before the stationary phase (OD_600nm_=1) by centrifugation. Then, cells were washed first with 2.0M LiAc and then 0.4M NaOH for 5 min on ice. Cells were finally boiled for five min at 95°C in Laemmli sample buffer (Bio-Rad, Hercules, CA, USA) supplemented with 100mM DTT.

Proteins were separated in 4-15% Mini-PROTEAN^®^ TGX Stain Free™ Protein gels (Bio-Rad, Hercules, CA, USA) and transferred to a nitrocellulose membrane (Hybond™-P, GE Healthcare, Chicago, IL, USA) using the semi-dry Trans-Blot^®^ Turbo™ Transfer System (Bio-Rad, Hercules, CA, USA). Membranes were blocked for an hour at room temperature with 5% milk powder diluted in PBS supplemented with 0.1% Tween-20 and then incubated with a primary antibody overnight at 4°C and one hour with a secondary horseradish peroxidase-conjugated antibody at room temperature. Membranes were developed with Enhanced Chemiluminescence ECL Plus™ (ThermoFisher Scientific, Waltham, MA, USA) and imaged with ImageQuant™ LAS 4000 (GE Healthcare, Chicago, IL, USA). The antibodies used in this study were polyclonal rabbit anti-human HINT1 antibody (1:1000, Sigma, San Luis, MO, USA), and to demonstrate equal loading mouse monoclonal anti-β-actin antibody (1:5000, Sigma, San Luis, MO, USA), mouse polyclonal anti-α-tubulin antibody (1:5000, Abcam, Cambridge, UK) or mouse polyclonal anti-PKG antibody (1:10.000, Sigma, San Luis, MO, USA).

### Dimerization reaction

Cells were harvested and lysed in 0.2 triethanolamine, pH=8.0, with 0.3 mg/ml of dimethyl adipimidate (DMP, ThermoFisher Scientific, Waltham, MA, USA) for 1 hour with vertical mixing. The lysates were cleared, and protein concentration was measured before immunoblotting.

### Yeast strain and transformation

*S. cerevisiae* strain BY8-5c (MATα *ura3-52 his3Δ200 trp1Δ901 lys2-801 suc2-Δ9 leu2-3,112 hnt1Δ::URA3*)[11] was kindly provided by Dr. Brenner, University of Iowa, USA. Yeast cells were cultured in rich (YP) medium supplemented with 2% glucose. The transformation of BY8-5c with the pAG415GPD expression plasmids carrying WT or variant *HINT1* was done using the LiAc/SS carrier DNA/PEG method[36]. Positive clones were selected in minimal medium without Leucine (SD-Leu) supplemented with 2% glucose.

### Spot assay

Pre-cultures of the different yeast clones were grown overnight in SD-Leu supplemented with glucose. Absorbance was measured and adjusted to an optical density of OD_600nm_=1. Serial dilutions of each culture were performed, and 5µl culture was spotted on SD-Leu agar plates supplemented with either 2% glucose or 2% galactose. Plates were incubated for three days at 39°C.

## RESULTS

### Systematic evaluation of NMAN-associated variants using pathogenicity predictors defines their pathogenic character

To select HINT1 variants to be investigated in this study, we curated articles published between 2012 and 2025 that reported genetic alterations of HINT1 in patients with peripheral neuropathies. A total of 29 *HINT1* variants were retrieved (Table 1), including 21 missense, five stop-gain (p.Gln9Ter, p.Ser61Ter, p.Gln62Ter, p.Gln106Ter, p.Trp123Ter) [7, 9, 37, 38], two small exonic deletions (p.Phe33Leufs*22, p.His51Phefs*18) [39, 40], one deletion affecting the splice-donor site of the canonical transcript (c.112-1delG)[41], and one complex chromosomal rearrangement causing start-loss (g. [131164372_131165305delins131165015_131165143inv; 131165362_131165385del]) [42]. In most of the patients, the age of onset of the neuropathy was in the first decade of life, even though cases with later onset (early 30s) have also been reported. Neuromyotonia, the hallmark sign of the disease, was reported in 73% of the affected individuals.

**Table 1:**
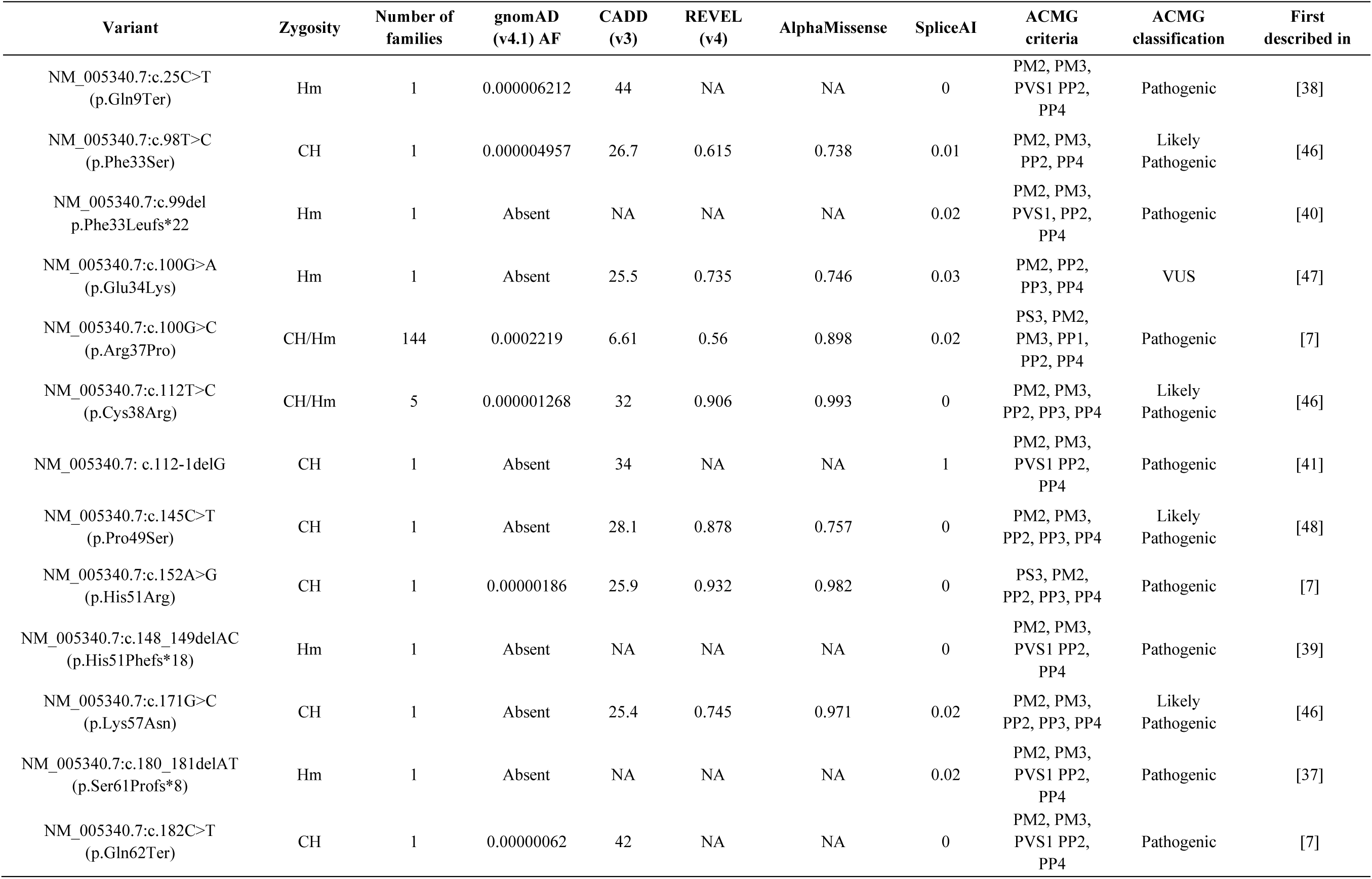
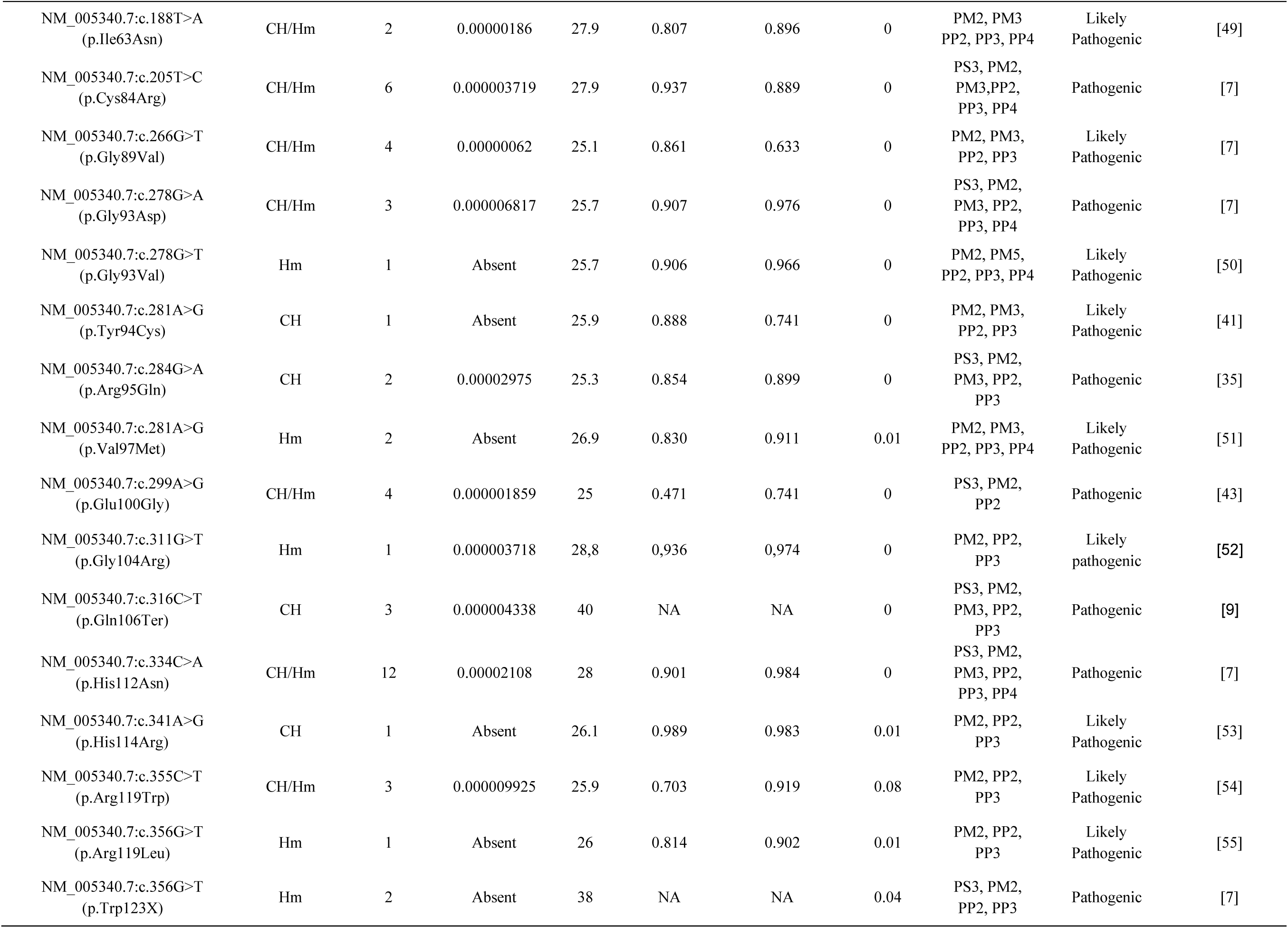
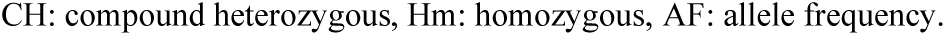
Summarized overview of variants identified in NMAN patients.

Overall, 12 out of 29 variants (41%) were recurrent and have been identified in more than two unrelated patients in either compound heterozygous or homozygous state. Among them, there were seven founder variants in Europe (p.Arg37Pro, p.Cys84Arg, p.Gly89Val, p.Arg95Gln, p.Glu100Gly, and p.His112Asn) [8, 35, 43] and Asia (p.Cys38Arg) [24]. Notably, the p.Arg37Pro allele was present in nearly 80% of all NMAN patients. The 17 remaining variants were identified in sporadic cases only: five were homozygous in non-consanguineous families, and 11 were in a compound heterozygous state (Table 1).

Most of the variants had a very low allele frequency (<0.0005%) across all represented genetic ancestries (gnomAD v 4.1), and a substantial fraction (15/28) was absent from population databases. According to *in silico* predictors of variant pathogenicity (including CADD [44], REVEL [24], and AlphaMissense [45]), 25/29 of the missense variants had a damaging effect. For p.Phe33Ser, p.Glu34Lys, p.Arg37Pro, p.Glu100Gly, there were conflicting results across the three predictors. We categorized all *HINT1* variants according to the American College of Medical Genetics and Genomics (ACMG) guidelines. One variant was classified with unknown significance (VUS) (p.Glu34Lys), thirteen were likely pathogenic, and seven were pathogenic. Two variants with conflicting *in silico* prediction scores (p.Arg37Pro and p.Glu100Gly) were classified as pathogenic by the ACMG guidelines based on the functional data available (PS3).

### ClinVar reported variants of uncertain significance in HINT1 highlight the need for re-assessment of pathogenicity

ClinVar is a public archive for human variation reporting variants found in patient samples, pathogenicity classifications, drug responses, submitter information, and additional supporting data. As such, it is a freely accessible resource for curating both published and unpublished data on variants observed in patients and has long been used for variant interpretation. A total of 121 HINT1 variants (excluding copy number variants impacting HINT1 together with multiple other genes) have been submitted to the ClinVar database to date (Additional File 1). Approximately 21.5% of them (26/121) have a germline classification of pathogenic or likely pathogenic. Two percent of the ClinVar variants (3/121) have conflicting interpretations of pathogenicity, and interestingly, these were all previously reported in NMAN cases (p.Ile63Asn, p.Arg95Gln, p.Arg119Trp). The remaining 38% of the variants (43/121) are classified as benign or likely benign. Additionally, 40% of the ClinVar variants (49/121) were reported to have uncertain significance, including three variants identified in NMAN patients (p.Tyr94Cys, p.Glu100Gly, p.His114Arg).

Using AlphaMissense, we calculated pathogenicity scores for all possible amino acid substitutions of each residue across the HINT1 sequence. By visualizing these scores in a heatmap, we identified regions that are either more tolerant to missense variants or serve as “hotspots” for substitutions with a deleterious effect(Fig. 1). We then integrated this data with the positions of reported NMAN-associated variants and those extracted from the ClinVar database. Our analysis revealed that most of the 21 missense variants analyzed in this study cluster within regions with high pathogenicity scores. Notably, the distribution of ClinVar variants with uncertain significance along the protein sequence differed significantly from that of the NMAN-associated variants (Fig. 1B), and they showed lower mean pathogenicity scores (Fig. 1C). This suggests that many VUS reported in the ClinVar database are likely benign, however experimental validation is necessary to clarify their true clinical significance.

**Figure 1.**
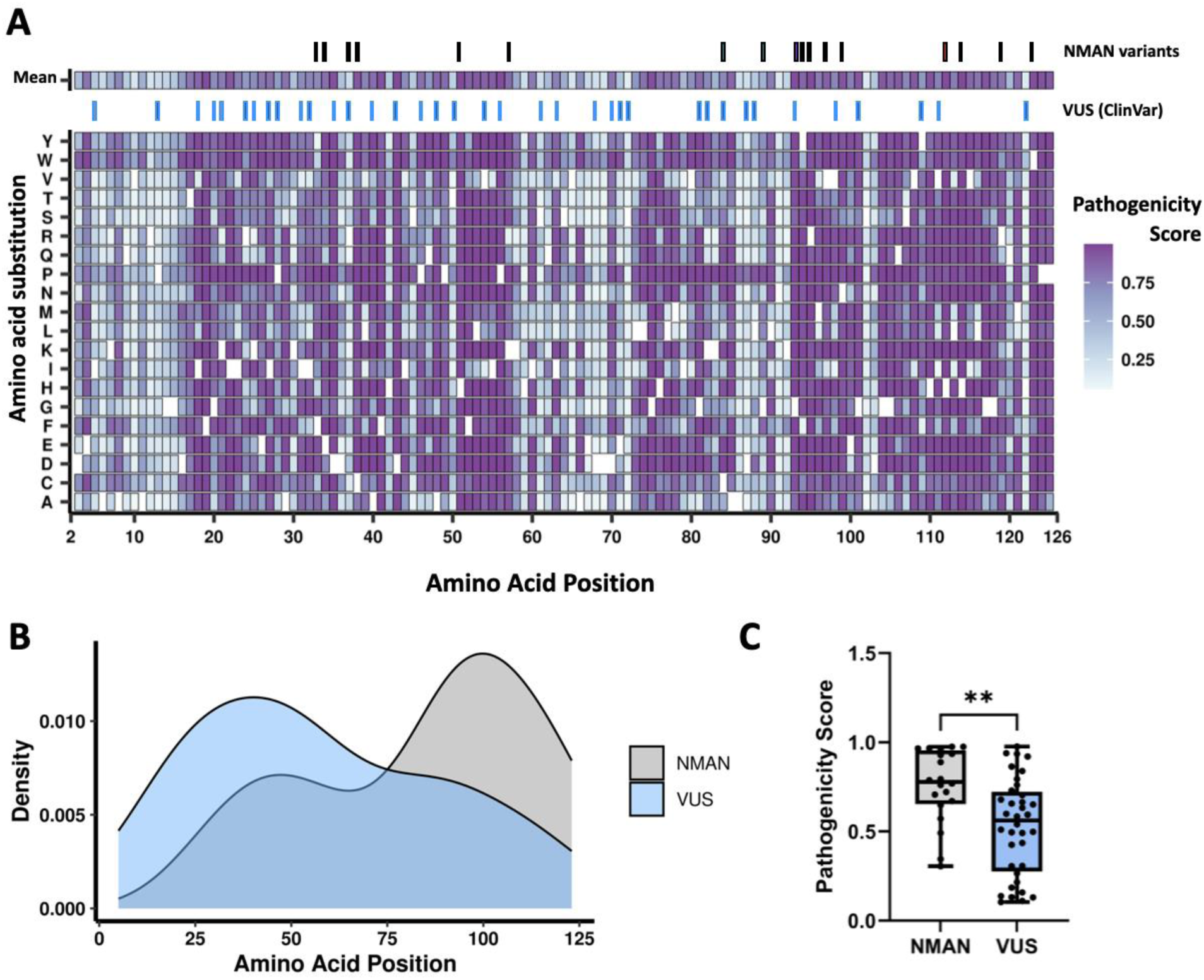
Pathogenicity scores for missense mutations across HINT1 primary structure calculated by AlphaMissense. **A)** Heat map representing every position in the HINT1 amino acid sequence and the pathogenicity score calculated by AlphaMissense of every possible amino acid substitution. Synonymous variants are depicted as white spaces. On top, the average pathogenicity score for each amino acid in the HINT1 sequence. Blue rectangles indicate the positions of the variants of uncertain significance (VUS) submitted to ClinVar. Black rectangles indicate the position of the variants reported in NMAN patients characterized in this study. **B)** Kernel density estimates were plotted to compare the distribution of the VUS reported in ClinVar (blue) and the NMAN-associated variants (grey). Wilcoxon Rank-sum test showed significant differences between the two distributions (p=0.004288). **C)** Comparison of the predicted pathogenicity score by AlphaMissense shows a significantly lower pathogenicity score (unpaired t-test, p=0.0016) for the VUS reported in ClinVar compared to the NMAN-associated variants

### A combination of computational approaches and 3D mapping of NMAN causal variants identifies structural clusters

To study the effect of each NMAN-associated variant on the HINT1 protein, we used a combinatorial computational approach where we assessed the impact of residue positions, solvent-accessible surface areas, protein stability, and conservation. Together, these allowed us to identify four (non-mutually exclusive) groups in which the variants showed commonalities. The first group contained all predicted loss-of-function (pLoF) genetic variants. These include four stop-gain variants (p.Gln9Ter, p.Ser61Ter, p.Gln62Ter, p.Gln106Ter), two small exonic deletions (p.Phe33Leufs*22, p.His51Phefs*18), the loss of splice donor-site of the canonical transcript (c.112-1delG), and the start-loss caused by a complex chromosomal rearrangement (g. [131164372_131165305delins131165015_131165143inv;131165362_131165385del]). In all cases, these variants either resulted in a truncated polypeptide lacking the characteristic HIT motif, potentially triggering nonsense-mediated mRNA decay, or prevented protein production due to start-loss. Therefore, they all lead to non-functional proteins and, hence, are classified as pathogenic (Fig. 2A, black). The remaining genetic variants were classified into three additional structural clusters based on their spatial distribution on the 3D protein structure and the kind of potential change they might impart on the protein. These include a) the catalytic pocket, b) the dimer interface, and c) the distal β-hairpin and nearby loop (Fig. 2A, B).

**Figure 2.**
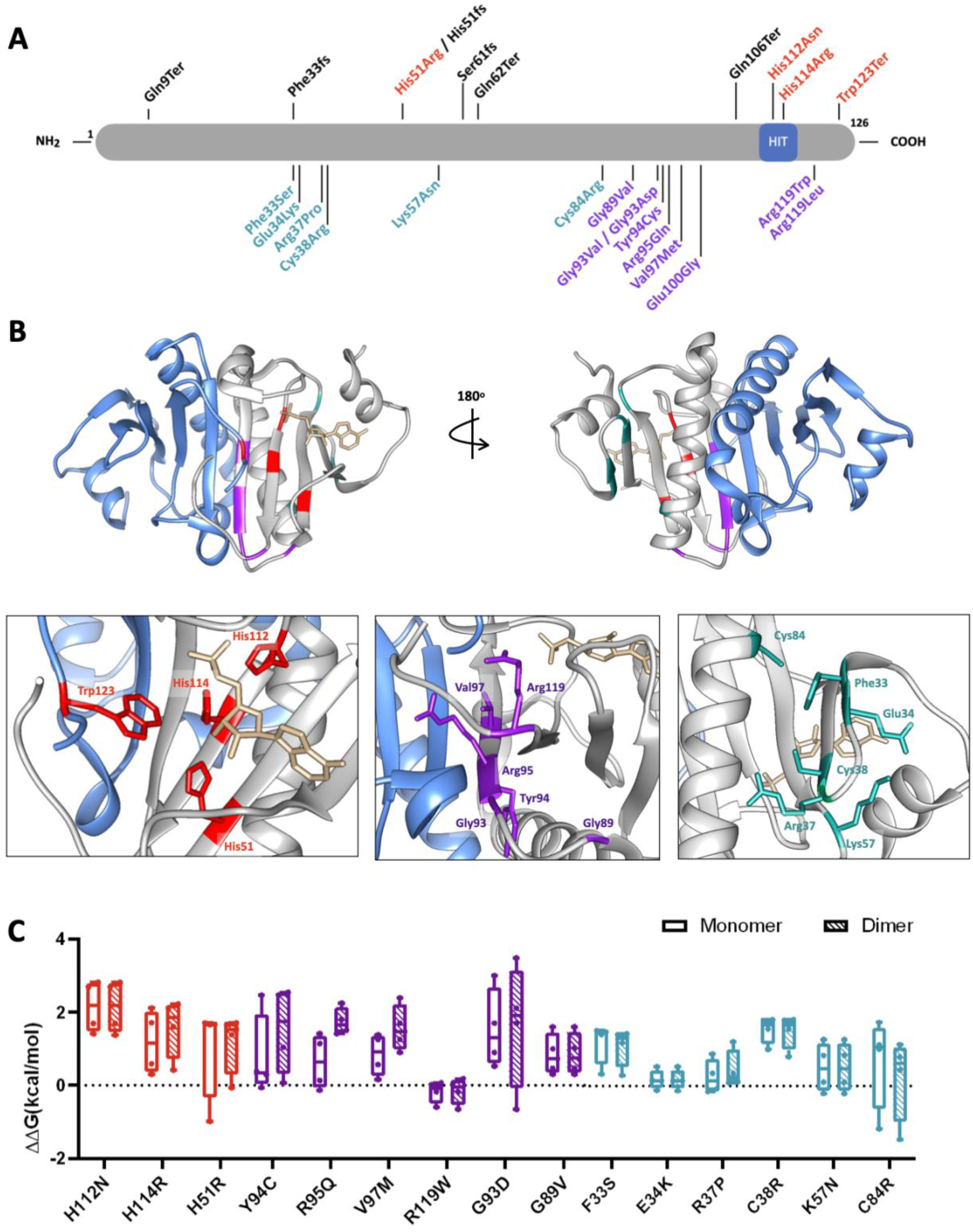
Classification of HINT1 missense variants in structural clusters. **A)** Mapping of the HINT1 missense variants onto the HINT1 protein sequence. The highly conserved HIT motif is represented as a blue box. **B)** Mapping of the residues affected by NMAN-causal variants in the crystal structure of the HINT1 dimer (PDB: 5KLZ). Each monomer is shown in “cartoon” representations and in different colors (blue and silver); AMP is indicated in gold using a “sticks” representation in the catalytic pocket of the enzyme. Close-ups of the three structural clusters identified are shown on the bottom panel. The variants are colored according to the cluster classification. Red: variants at the catalytic site. Purple: variants at the dimer interface. Green: variants in the distal β-hairpin and the hairpin-helix region. **C)** Predicted destabilization effects of the NMAN missense variants on the HINT1 dimer and monomer stability, respectively. PDB structure 5KLZ chain A+B was used as input for evaluating the effects on dimer stability, and chain B alone for monomer stability. The ΔΔG distributions represent the ΔΔG values obtained using the different stability predictors (*see Materials & Methods for details*). Values calculated by FoldX showed the most significant fluctuations, often differing from those of other methods, and were therefore excluded from the plot (for ΔΔG values computed by all methods, see Additional File 3). Because p.Trp123Ter is not a missense variant, the destabilizing effect of early termination of the protein sequence could not be calculated by the stability predictors.

The catalytic pocket cluster (Fig. 2A, B, red) includes all residues directly involved in the phosphoramidase activity of HINT1. These residues are either part of the highly conserved HIT motif (p.His112, p.His114), they help stabilize the product intermediates (p.His51), or contribute to substrate recognition (p.Trp123). Additionally, two residues (p.His114, p.Trp123) contribute to the formation of the dimeric holoenzyme. The high conservation of the affected residues across species highlights their essential function and suggests that their substitution will have a detrimental effect on the enzymatic activity (Additional File 2).

Accordingly, all residue substitutions are predicted to destabilize both the monomeric and dimeric forms of HINT1 equally (Fig. 2C).

Many residues hit by NMAN missense substitutions (p.Gly89, p.Gly93, p.Tyr94, p.Arg95, p.Val97) cluster at the dimer interface in an antiparallel β-sheet (Fig. 2A, B, purple). Additionally, the p.Arg119 residue, although it does not reside at the dimer interface, forms hydrogen bonds with the C-terminus of the neighboring monomer, further stabilizing the contacts between the two monomers. The majority of the surface area of the residues in this cluster is buried upon formation of the HINT1 wild-type (hWT) dimer (*see Methods for details*), and significantly more than any of the other variants analyzed (BSA_R_ in Additional File 3), confirming their essential contribution to the stability of the dimer interface. Their substitution with polar or bulkier residues is expected to disrupt the dimer interface, thereby destabilizing the protein. This destabilization is predicted to be higher for the dimeric form compared to the monomeric form, reinforcing their role in dimer formation (Fig. 2C). The residues of this cluster are conserved across species, with the highest conservation seen for p.Tyr94 and p.Arg119 (Additional File 2).

The third cluster includes variants affecting residues lying far from the dimer interface and the catalytic pocket. They are located at the surface of the protein, featuring the smallest fraction of buried surface area, and hence remain essentially accessible to solvent in both the monomeric and dimeric forms (Additional File 3). Four residues (p.Phe33, p.Glu34, p.Arg37, p.Cys38) are positioned in a β-hairpin at the N-terminus. The p.Lys57 residue is adjacent to the β-hairpin, and the p.Cys84 is in an α-helix that packs against the β-sheet formed by the hairpin. This cluster contributes to preserving the native helix-sheet interactions (Fig. 2, green). Interestingly, this part of the protein exhibits the lowest degree of homology among species, with significant conservation observed only across mammals (Additional File 2). The p.Phe33Ser, p.Glu34Lys, p.Arg37Pro, p.Lys57Asn, and p.Cys84Arg variants of the β-hairpin cluster are predicted to have only mild destabilizing effects on the dimer and monomer, while a higher destabilizing effect is predicted for the p.Cys38Arg variant (Fig. 2C).

### Experimental characterization using functional assays reveals commonalities among variants within the same structural cluster

To functionally assess the impact of the missense variants on the HINT1 structure, we transiently overexpressed the disease-causing alleles in a *HINT1* knock-out (KO) HeLa cell line[35] and quantified the resulting exogenous HINT1 protein abundance by immunoblotting (Fig. 2A). We modeled the most recurrent substitutions for p.Gly93 and p.Arg119 (p.Gly93Asp and p.Arg119Arg) even though other alterations of the same residues are also reported (p.Gly93Val and p.Arg119Leu). When disease-causing alleles impacting the catalytic pocket were transiently expressed in HeLa cells, no HINT1 protein was detected by immunoblotting. The p.His112Asn allele was a notable exception to this, with the resulting HINT1 protein expressed at comparable levels to the endogenous WT protein (Fig. 3A). Our finding aligns well with previous reports showing similar stability and structure of this variant compared to the hWT protein. The six substitutions of the dimer interface cluster had different impacts on the overall stability of the protein in HeLa cells (Fig. 3A). Two of them (p.Val97Met, p.Arg119Trp) heavily impacted HINT1 stability, and the resulting proteins could not be detected. Two others (p.Gly93Asp, p.Arg95Gln) caused only a reduction in protein expression levels, whereas the last two variants (p.Gly89Val and p.Tyr94Cys) resulted in proteins expressed at levels similar to that of the hWT protein. Lastly, the β-hairpin cluster variants resulted in proteins with high instability. Only the p.Phe33Ser allele was expressed at similar levels as the hWT protein, whereas only faint traces of the p.Lys57Asn and p.Cys84Arg proteins were detectable by immunoblotting (Fig. 3A).

**Figure 3:**
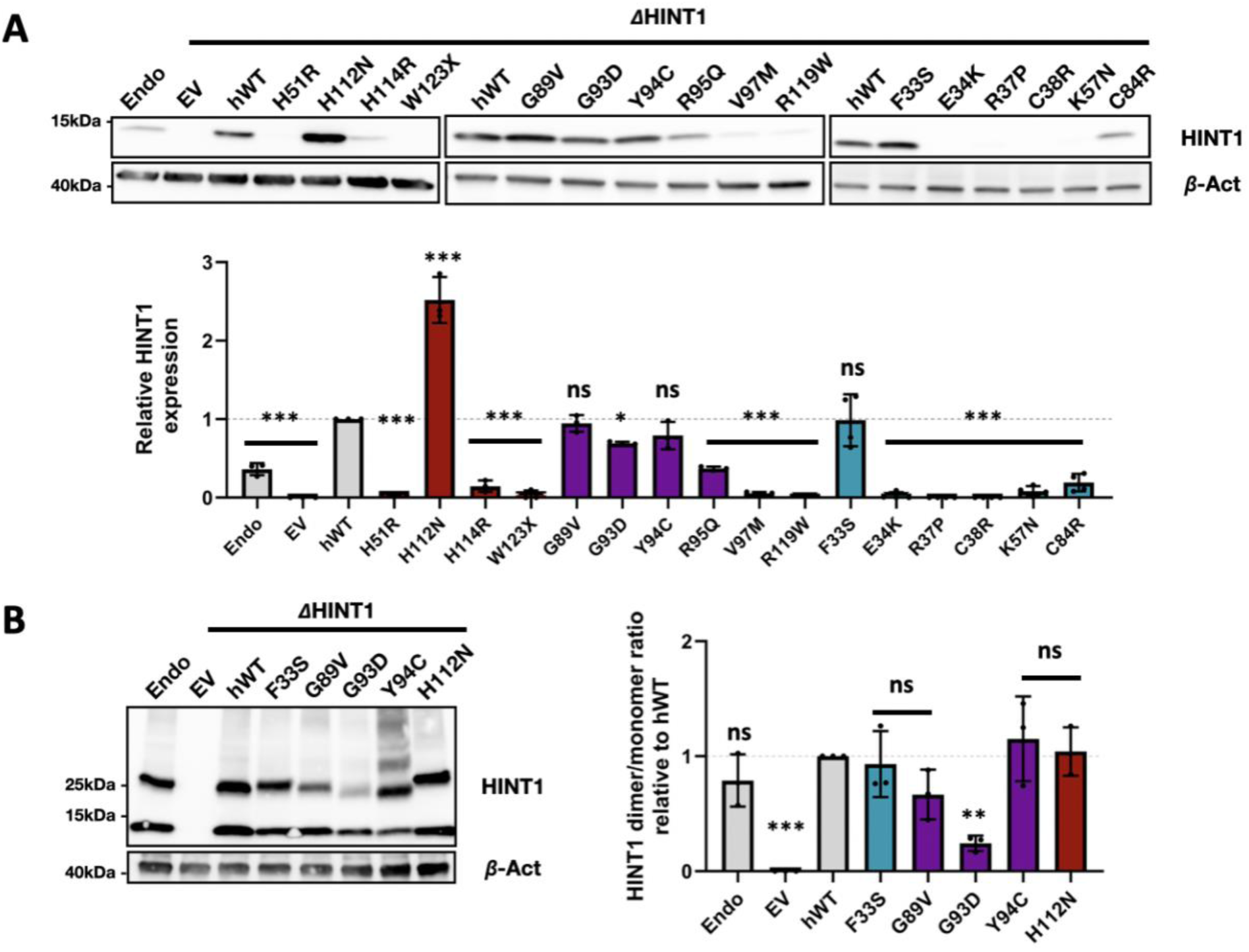
In cellulo characterization of NMAN-associated missense variants in HINT1. **(A)** Expression constructs carrying an NMAN allele were transiently expressed in HeLa *HINT1* KO cells, and the HINT1 protein levels were assessed by semiquantitative immunoblotting. Equal loading was validated with β-actin, and relative HINT1 expression was normalized to hWT expression. The graph represents the relative quantification of band intensities for three independent replicates. **(B)** Immunoblotting after dimerization reaction using DMP. HINT1 protein expression levels in monomeric (14 kDa) and dimeric (28 kDa) forms could be detected. The level of dimer formation was quantified relative to the monomer expression and normalized to the hWT. One-way ANOVA with Dunnet multiple comparison tests was used in all western-blot quantifications. Bar charts are presented as means with standard error of the mean (s.e.m). ns = no significant, *p<0.05, **p<0.01, *** p<0.001

Since HINT1 enzymatic function relies on the correct formation of HINT1 homodimers, we investigated if HINT1 dimerization was impaired in those alleles that did not cause protein instability in our previous analyses. To this end, we treated the cell lysates with dimethyl pimelimidate (DMP), a cross-linking reagent active towards primary amines. DMP has a short linker (7 Å), allowing the study of protein quaternary structures by covalently binding the different subunits together. Protein levels of the monomeric and the dimeric forms of HINT1 were detected via immunoblotting (Fig. 3B), and the ability to form dimers was quantified as a ratio of HINT1 dimer over the HINT1 monomer normalized to the HINT1-hWT overexpression ratio. In cells expressing hWT proteins, two bands were detected, one at 14 kDa (monomer) and another at 28 kDa (dimer). The tested altered proteins showed a heterogenous effect on dimer formation efficiency (Fig. 3B). Only the p.Gly93Asp protein remained monomeric and showed severe dimerization impairment. The remaining proteins were able to form HINT1 dimers, similar to the hWT protein; however, they showed additional features. The p.Phe33Ser showed a slight shift in the molecular weight of the dimer. An even larger shift was observed for the p.His112Asn protein, suggesting the likely presence of an additional binding partner. The p.Tyr94Cys protein size-separated into several bands, indicating that it is prone to aggregation or interacts with another partner(s). Finally, only the p.Gly89Val protein could form dimers as the hWT protein.

Currently, the only *in vivo* model for HINT1 neuropathy uses a yeast strain deficient for *HNT1* (HINT1 orthologue). This strain exhibits a growth deficit under restrictive conditions (Galactose, 39°C) that can be complemented by the human HINT1 wild-type protein (hWT), thus providing an easy readout for testing HINT1 functionality[7]. We used this model to assess the detrimental effect of the HINT1 missense variants by first checking the stability of the proteins in the yeast by immunoblotting, followed by testing their functionality based on their ability to rescue the yeast growth deficit.

All altered proteins from the catalytic cluster were stably expressed in yeast with only p.His51Arg and, to a lesser extent, p.Trp123Ter, showing relatively lower expression levels (Fig. 4A). Despite their presence, none of them could rescue the growth deficiency, suggesting a loss of catalytic function (Fig. 4B). The proteins harboring variants at dimer interface were all expressed at similar levels to the WT protein, except for p.Val97Met, whose expression was slightly reduced (Fig. 4A). Despite all the corresponding proteins being stably expressed in yeast, only p.Gly89Val caused a complete rescue of the HINT1 growth phenotype, whereas p.Tyr94Cys and p.Arg119Trp enabled only partial recovery of the yeast growth under stress conditions. The remaining three substitutions (p.Gly93Asp, p.Arg95Gln, p.Val97Met) could not detectably restore yeast growth, confirming their loss of function (Fig. 4B). The higher protein instability caused by the substitutions of the β-hairpin residues was also apparent in yeast except for the p.Glu34Lys allele, which, although highly unstable in HeLa cells, was stably expressed in yeast (Fig. 4B). Additionally, the p.Cys84Arg and p.Lys57Asn proteins were expressed at lower levels than WT. In contrast, the remaining proteins (p.Arg37Pro and p.Cys38Arg) exhibited high instability, with minimal detectable expression. Regarding the complementation of the yeast growth deficit, only the p.Phe33Ser and p.Glu34Lys were able to fully restore the growth as efficiently as the hWT protein (Fig. 4C), while the p.Cys84Arg and p.Lys57Asn could only do so partially.

**Figure 4.**
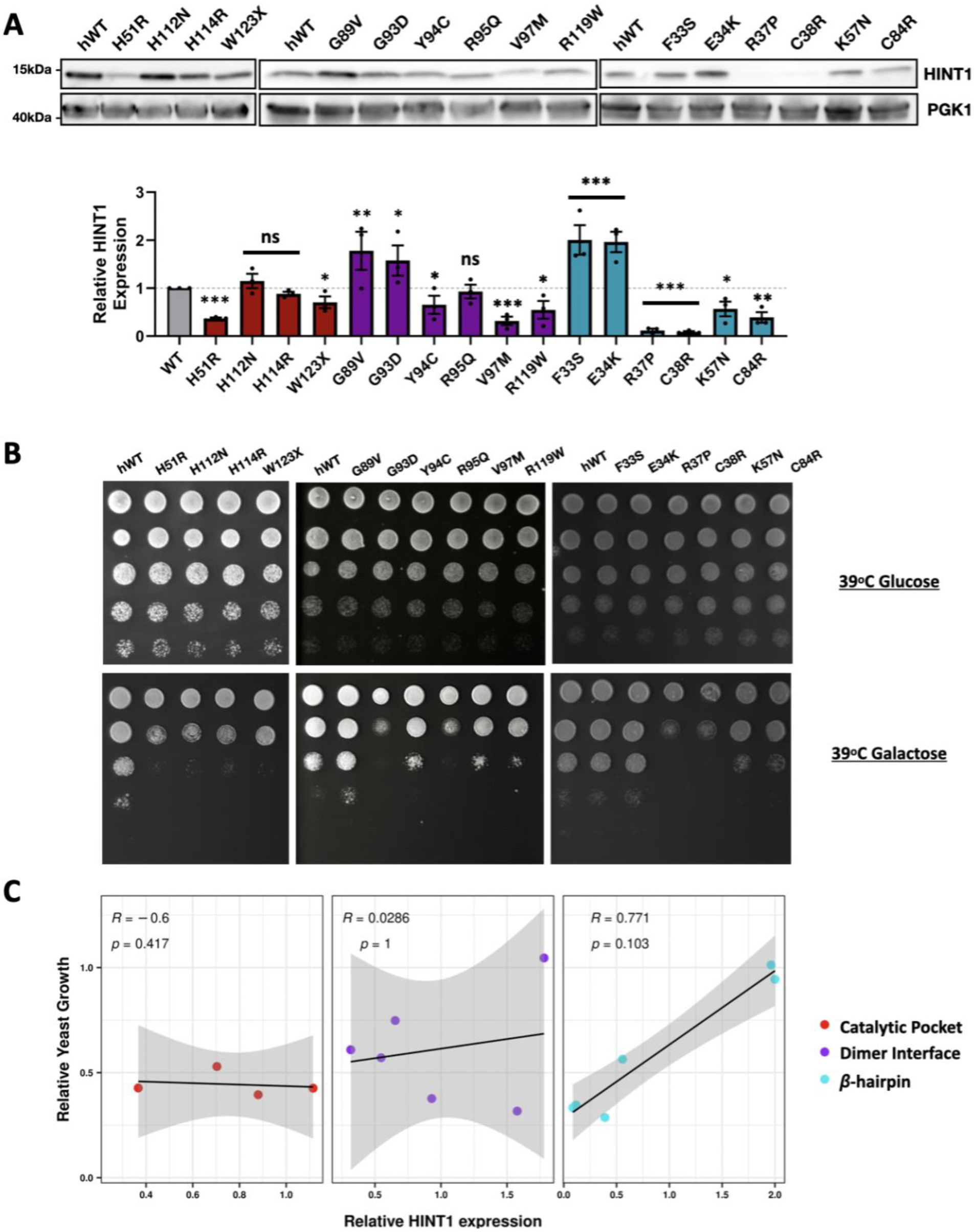
In vivo characterization of NMAN-causing variants. **A)** Immunoblotting analysis of total protein extracts of a yeast HNT1-KO strain expressing the human mutant or WT HINT1 protein. Relative quantification of HINT1 levels was normalized to the expression of hWT protein. **B)** Genetic complementation analysis in a *HNT1*-deficient yeast strain. A serial dilution of yeast culture was spotted in a plate and let grow for three days under permissive (2% Glucose, 39°C) and stress (2% Galactose, 39°C) conditions. **C)** Linear coefficient correlations between the HINT1 protein expression and yeast growth relative to the hWT conditions. The correlations are calculated cluster-wise and Spearman’s correlation coefficients and corresponding p-values are indicated for each cluster.

We assessed the relationship between HINT1 protein expression and their ability to rescue the yeast growth by calculating the Spearman’s correlation coefficient between those two variables for the three variant clusters (Fig. 4C). Only the modified proteins of the β-hairpin cluster showed a strong positive linear correlation between the expression levels of the altered proteins and their ability to rescue the growth phenotype in the yeast. Although the correlation was not significant, likely due to the small sample size, this suggests that these amino acid substitutions do not detrimentally impact the HINT1 enzymatic activity.

## DISCUSSION

The continuous effort to discover disease-causing genes is crucial for improving diagnostic accuracy, as well as for advancing therapy development, since understanding the genetic basis enables the identification of novel drug targets and facilitates precision medicine efforts. Next-generation sequencing (NGS) technologies have dramatically increased the discovery of novel variants likely causing genetic diseases; however, this larger number of identified variants (mostly missense) brings along the challenge of clinically interpreting their pathogenicity. Strategies to infer variant causality include co-segregation with disease phenotype, identification of additional affected families carrying the same variant, low allele frequency in controls, computational predictions suggestive of pathogenicity, and *in vitro* and/or *in vivo* experimental evidence supporting pathogenicity [1, 3, 56]. Variant interpretation, especially in ultra-rare disorders, relies on robust computational and functional characterization strategies.

In this study, we used NMAN as an exemplary ultra-rare disease where the causal relationship with HINT1 is well-established and variant interpretation is facilitated by the presence of the clinical hallmark of neuromyotonia. Nearly 30 variants spread along the protein sequence are reported in patients with peripheral neuropathy, yet functional data are available for only approximately one-fourth of them. Strikingly, approximately 40% of the *HINT1* variants reported in the ClinVar database have a germline classification of uncertain significance, highlighting the need for re-assessment of their clinical significance. Among the criteria given in the guidelines for variant interpretation by ACMG guidelines[57], one of the most widely applicable parameters is *in silico* pathogenicity prediction. Many *in silico* tools exist using different algorithms for evaluating variant impact; however, achieving concordance among them is an increasing challenge[58]. We used three main *in silico* meta-predictors to assess the pathogenicity of multiple missense variants in *HINT1* (CADD, REVEL, and AlphaMissense) and often also observed conflicting interpretations (see Table 1). Among other factors, the function of proteins relies on their intact native 3D conformation. However, none of the available pathogenicity predictors considers the quaternary structure of the proteins [59]. This is especially important for HINT1 since its physiologically active state in the cell is as a homodimer. Therefore, we additionally calculated the effect of the variants on the monomer and dimer stability (folding free energies) using six different algorithms (Fig. 2) and the change of solvent accessibility area upon homodimer formation (Additional File 3). Taken together, these analyses allowed us to infer the contribution of each residue on dimer formation and how it was impacted by the amino acid substitution.

Based on our combined approach, we could classify the HINT1 variants into four different groups: *1)* truncation variants that lead to lack of HINT1 expression (i.e., p.Ser61Ter), *2)* variants that render a dysfunctional catalytic pocket (i.e., p.His112Asn), *3)* variants that impact the dimer interface (i.e. p.Gly93Asp), and *4)* variants that affect the distal β-hairpin and nearby loop (i.e. p.Arg37Pro). From a functional perspective, overlaps exist between the variants of the dimer interface and the catalytic pocket. Since HINT1 enzymatic cleft is formed by the two monomers, it is not surprising that residues of the catalytic pocket (p.His114 and p.Trp123) are involved in maintaining the structure of the dimer or alternatively, variants destabilizing the dimeric HINT1 form (p.Gly93Asp) completely abolish the enzymatic activity. Notably, variants targeting residues located in the β-hairpin appear to have the most significant destabilizing effect. However, this destabilization effect was not as pronounced in the yeast allowing us to test the functionality of mutant proteins that are otherwise degraded (Fig. 4). Importantly, we showed that they are unstable but enzymatically active, and their activity correlates with the HINT1 expression levels. Similarly, previous *in vitro* studies showed that the p.Cys84Arg allele mildly affected the HINT1 3D structure and stability but retained its enzymatic activity despite being degraded in patient-derived cells[7, 60].

Our characterization of each of the NMAN-associated proteins *in cellulo* and *in vivo* provided with functional evidence of pathogenicity for additional seven variants classified as likely pathogenic (p.Cys38Arg, p.Lys57Asn, p.Tyr94Cys, p.Val97Met, p.His114Arg, p.Arg119Trp) or VUS (p.Glu34Lys) by the ACMG guidelines. Moreover, the functional assays demonstrated the deleterious character of the variants where the *in silico* pathogenicity predictors showed contradictory results (i.e., p.Glu34Lys). Studying the effect of the HINT1 missense variants on the protein function also allowed us to gain mechanistic insights into NMAN pathomechanism. For instance, most missense variants led to the expression of an unstable protein in patients, effectively leading to loss of protein function. Notably, HINT1 is a versatile protein with different properties within the cell, and it is unclear which of the activities is associated with NMAN. The functional characterization we performed suggests that the enzymatic activity of HINT1 is essential in the disease process. For example, the p.His112Asn allele results in a stable but enzymatically inactive protein, and therefore, it is unable to rescue the growth deficiency in *HNT1*-KO yeast. However, this altered protein has been demonstrated to still function as a transcriptional regulator of the p53 tumor suppressor gene in cancerous cells. In addition, two substitutions (p.Phe33Ser and p.Gly89Val) showed comparable functionality to the hWT protein in all our assays. Despite appearing benign in these tests, both genetic and clinical evidence strongly support their pathogenic nature [7]. In fact, *in vitro,* these substitutions have been shown to disrupt HINT1 SUMOylase activity [14] and their binding with several interacting partners, such as calmodulin or the mu-opioid receptor, among others [61]. These variants are, therefore, of particular interest in further uncovering both known and unknown HINT1 functions that might contribute to disease pathology.

Taken together, all NMAN-causal variants lead to a similar outcome, i.e., lack of a functional HINT1 enzyme in the cell. However, the establishment of four different classes of pathogenic variants based on their structural and molecular characteristics suggests that a “one-size-fits-all" treatment would not apply to all NMAN patients. The assay pipeline we developed can be used not only to assess the pathogenicity of any new and uncharacterized variant in HINT1 but also serves as a tool to define the variant sub-class to tailor the therapeutic strategy accordingly. For example, the NMAN-associated variants belonging to the β-hairpin cluster preserve HINT1 enzymatic activity and show a linear correlation between HINT1 protein expression and the rescue of yeast growth. Notably, most of the recurrent and high-frequency variants (e.g., p.Arg37Pro) belong to this cluster. Therefore, they seem to be ideal candidates for a pharmacochaperone therapeutic approach. This approach involves a small molecule that could bind and stabilize the mutant HINT1 protein *in vivo* and in this way restore its functionality. A similar strategy has been successfully applied to treat cystic fibrosis, where a small molecule stabilizes the cystic fibrosis transmembrane conductance regulator (CFTR) channel, reversing the effects of the pathogenic variant and slowing the disease progression [62].

## CONCLUSIONS

In summary, our standardized experimental pipeline allows for the characterization of newly discovered HINT1 variants in the context of NMAN. Our work highlights the value of combining structural and functional approaches to understand better the underlying disease mechanisms. Our findings provide the rational basis for patient stratification and development of personalized treatment strategies.

## LIST OF ABBREVIATIONS

ACMG: American College of Medical Genetics and Genomics
AF: Allele frequency
CADD: Combined Annotation Dependent Depletion
CH: Compound heterozygous
CMT: Charcot-Marie-Tooth disease
DMP: Dimethyl pimelimidat
FDA: U.S. Food and Drug Administration
gnomAD: Genome Aggregation Database
Hm: Homozygous
hWT: human HINT1 wild-type
kDA: kilodalton
KO: Knock-out
NGS: Next-generation sequencing
NMAN: Neuromyotonia-associated Axonal Neuropathy
PDB: Protein Data Bank
REVEL: Rare Exome Variant Ensemble Learner
VUS: Variant of Uncertain Significance

## SUPPLEMENTARY INFORMATION

The following supplementary material is available:

**Additional File 1:** Overview of all HINT1 variants submitted to the ClinVar database.

**Additional File 2:** NMAN-associated missense variants mapped to the ungapped multiple sequence alignment of HINT1.

**Additional File 3:** Overview of the ΔΔG distributions calculated using different stability predictors and the solvent-accessible surface area.

## DECLARATIONS

## Ethics approval and consent to participate

Not applicable

## Consent for publication

Not applicable

## Availability of data and materials

All data generated or analyzed during this study are included in this published article and its supplementary information files

## Authors’ contributions

SAB, TL, KP, SW, AJ: conception and design of the study; SAB, TL, AC, LLPR, KP, AJ, SW: acquisition and analysis of the data; SAB, TL: data visualization; SAB, TL, AC, SW, AJ: drafting the text. All authors read and approved the final manuscript.

## Acknowledgements

Not applicable

