## Additional File 2 for "Structural clustering and functional profiling of NMAN-causing variants in HINT1 suggest personalized therapeutic strategies"

**Supplementary Figures**

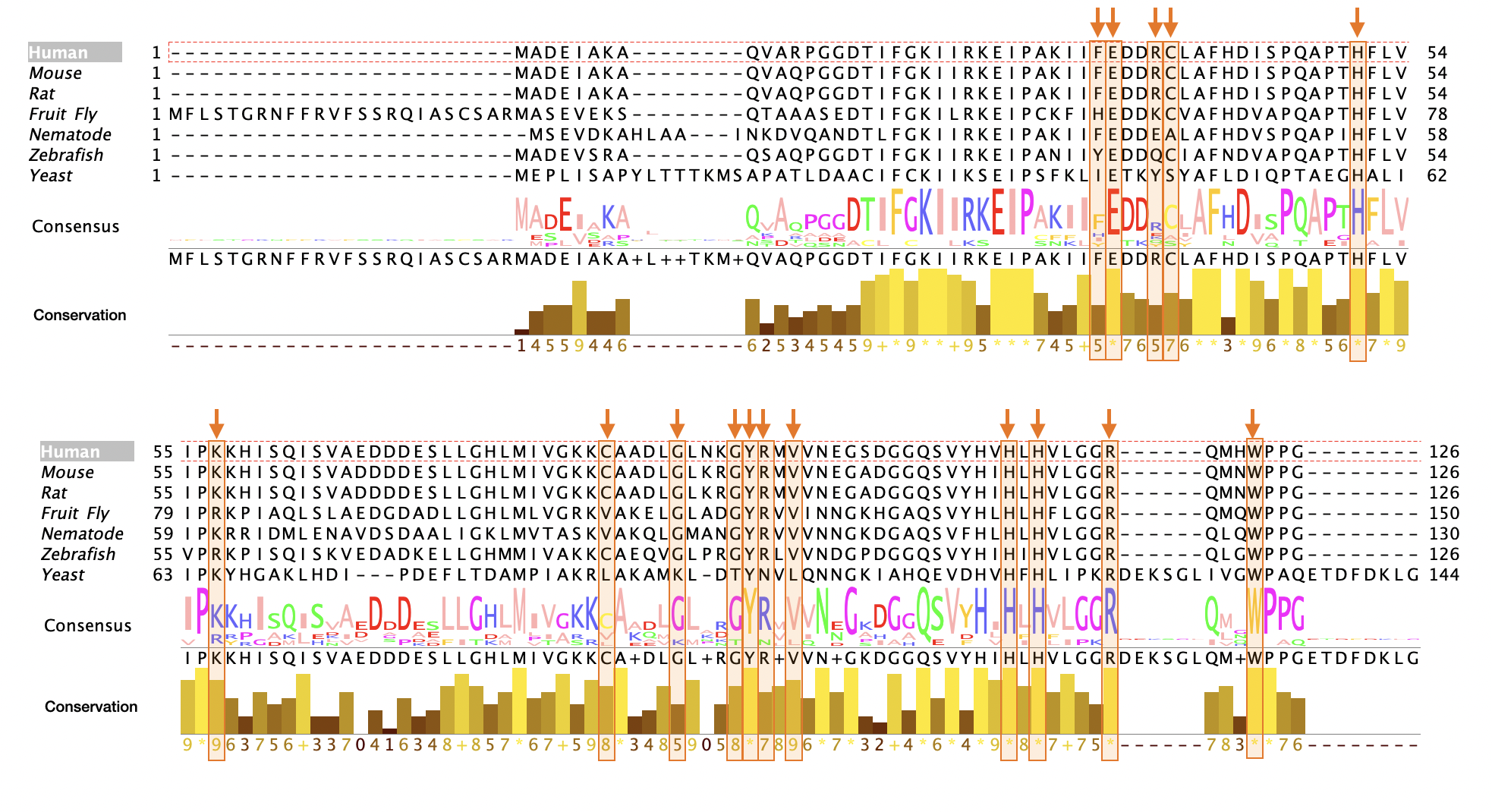

**Supplementary Figure 1. NMAN-associated missense variants mapped to the ungapped multiple sequence alignment of HINT1.**

The multiple sequence alignment of eukaryotic HINT1 orthologs from the OMA database aligned by ClustalOmega. Longer sequences than the human HINT1 sequence (highlighted on top) are hidden. Residues affected by NMAN mutations are indicated with red arrows above the alignment. Conservation is calculated by the AMAS method^1^ implemented in Jalview^2^. Quality reflects the normalized measure of the sum of pairwise ratios of the BLOSUM62 scores for all mutations and each residue’s conserved BLOSUM62 score. The histogram above the consensus displays the identity percentage of the column with the consensus residue.
